# Retinal image motion distracts visual motion memory even when generated by eye movement

**DOI:** 10.1101/2025.05.07.652578

**Authors:** Takeshi Miyamoto, Kosuke Numasawa

## Abstract

Interference between visual short-term memory (VSTM) and task-irrelevant sensory distractors is a well-documented phenomenon across a wide range of visual features, which is believed to stem from neural interactions between mnemonic and sensory processing. An overlooked question in the ongoing debate is whether this VSTM distraction is linked to the original sensory information or the ultimate perceptual experience. Here we addressed this issue by leveraging the perceptual invariance during ocular tracking of an object (i.e., smooth pursuit), where retinal image motion induced by smooth pursuit is encoded in early visual areas involved in motion processing, including the middle temporal (MT) and the medial superior temporal (MST) areas, but is perceptually suppressed. Our results showed that retinal image motion during the VSTM maintenance (delay period) attracted the memorized motion speed, consistent with previous studies. Importantly, the impact was comparable whether the retinal image motion was due to physical displacement of objects in world coordinates or to apparent motion induced by smooth pursuit without actual motion in world coordinates. In fact, observers’ responses in the VSTM task were partially predicted by retinal image motion during the delay period by a cross-condition classifier, where the distraction effect induced by the smooth pursuit-induced apparent motion was replicable using a classifier trained on data from the condition with the world-coordinate motion, and vice versa. These findings provide behavioral evidence that sensory inputs can distract VSTM without conscious perception and also suggest that the VSTM system shares neural substrates with sensory processing, but not with perception.

**Significance Statement:** Visual short-term memory (VSTM) is a temporary store for visual features and plays a crucial role in cognitive behaviors such as attention allocation and decision-making. VSTM is often distracted by task-irrelevant visual stimuli during its maintenance, particularly when memoranda and distractors are highly similar. In this study, we used a VSTM task for motion and demonstrated that VSTM can be distracted even when observers are almost not perceiving the distractor. This interference by perceptually suppressed sensory distractors may stem from the independence of perceptual invariance systems and those responsible for encoding VSTM based on sensory inputs.

## Introduction

Visual short-term memory (VSTM), also referred to as visual working memory (VWM), is the ability to retain and utilize visual information for goal-directed behaviors over short durations, even after the physical signals are no longer present on the retina (1). This ability underpins a wide range of cognitive functions and behaviors (2, 3). To function effectively, VSTM must resist interference by new sensory inputs that are irrelevant to the task at hand. From this perspective, the ability to maintain stable memory representations in the face of sensory distractors is considered a key characteristic of individual’s VSTM capacity (4, 5). A growing body of evidence from psychophysical, pharmacological, and neurophysiological studies has established that memorized visual information is robustly protected against distractors through various mechanisms related to cognitive control processes (reviewed in (6)). However, it is also known that distractors presented between the sample and recall periods (hereinafter referred to as the delay period) can induce a bias that subtly attracts the memory representation along a task-relevant feature dimension (6). This attractive bias is consistent in VSTM of diverse visual features and is most pronounced when memoranda and distractors are highly similar (7–9). This characteristic of VSTM has often been interpreted as evidence for the sensory recruitment hypothesis, which posits that sensory and memory processing share neural substrates (10).

Indeed, functional magnetic resonance imaging (fMRI) studies have successfully decoded VSTM contents from the blood-oxygen-level-dependent (BOLD) signals in early visual areas during the delay period (11, 12). Furthermore, studies with single-unit recordings have demonstrated that the prefrontal cortex (PFC), which plays a central role in the working memory network, contains neurons that encode either ongoing sensory inputs or memory, or both (13, 14). Although the specific brain region involved in the sensory-memory interference remains a topic of debate, the distractor-induced bias in VSTM likely originates from neural interactions in the region including sensory and mnemonic representations.

An overlooked but important question is whether the VSTM distraction is linked to original sensory signals or with ultimate perception. Perceptual experience of the world does not always faithfully reflect sensory inputs, yet this distinction has not been adequately explored in the context of VSTM. For example, when observers track a moving target with their gaze (i.e., smooth pursuit eye movement), the visual image of stationary background objects moves in the opposite direction on the retina, yet this motion is not consciously perceived. This perceptual invariance during smooth pursuit is achieved through the integration of retinal and extraretinal signals within the circuit of processing visual motion including the dorsal stream (15, 16). Note that the system for perceptual invariance does not suppress the encoding of retinal signals during smooth pursuit. Neurons in the middle temporal (MT) area, an early stage of the dorsal stream that receives inputs from direction-selective neurons in the primary visual cortex (V1) (17), encode retinal image motion even during smooth pursuit (18–20). In contrast, some neurons in the medial superior temporal (MST) area, where both retinal and extraretinal signals converge (21), compensate for pursuit-induced retinal motion, thereby providing motion information in screen (or world) coordinates (18–20). Such processing along the hierarchy of the dorsal stream is also evident in sensory and mnemonic representations. A study using single-unit recordings has shown that MT neurons predominantly encode information during the sample period, whereas MST neurons encode information during both the sample and delay periods though the proportion of delay-period encoding is smaller (22). Additionally, the PFC contains a similar proportion of neurons encoding information during the sample and delay periods (22).

Their further study has demonstrated that sensory and mnemonic information are partially encoded by distinct neuronal populations in the PFC, allowing for a linear neural population decoder to predict the direction of represented motion and determine whether it pertains to ongoing or memorized information (13). They concluded that the ongoing sensory representation reflects copies of information inherited from upstream sensory areas, and the mnemonic representation emerges as a result of local processing in the PFC (23). However, it remains unclear whether the inherited sensory signals involved in memory representations are identical to ultimate perception after compensation for eye movement. If they are not identical, even the apparent retinal image motion induced by smooth pursuit may act as a distractor for visual motion memory, despite being perceptually suppressed.

This question is somewhat analogous to whether unconscious (perceptually subliminal) sensory inputs can influence VSTM. Some studies in VSTM for object orientation have explored this issue, which suggest that potential distractors in VSTM are limited to consciously perceived sensory inputs. For example, a study using supraliminal and subliminal distractors has shown that encoding the orientation of an object was biased by a supraliminal distractor (i.e., one that is consciously perceived due to adequate presentation duration) but not by a subliminal distractor (i.e., one that is not consciously perceived due to its brief presentation duration) (24). Furthermore, another study using binocular rivalry has shown that recalling object orientation was not distorted when a delay-period distractor was presented to the non-dominant eye (i.e., the side where perception is suppressed) and simultaneously a mask is presented in the dominant eye (i.e., the perceived side) (25). These studies provided important insights through unique experimental designs; however, it is important to note that direct comparisons were challenging between the conscious and unconscious conditions in these studies because the unconscious conditions typically involved smaller or noisier sensory inputs compared to the conscious conditions. If these experimental stimuli suppressed the encoding of sensory inputs themselves (e.g., suppression of sensory representations in the non-dominant eye during binocular rivalry (26)), the distractors would not affect VSTM, regardless of whether VSTM is linked to original sensory signals or ultimate perception.

In the present study, we examined whether the distraction effect of retinal image motion during the delay period persists when the motion is induced by smooth pursuit. In our experiment, observers were asked to memorize the speed of visual motion and compare it with that of another motion. During the delay period, task-irrelevant motion was presented as a sensory distractor, consisting of either actual motion in world coordinates or apparent motion induced by smooth pursuit. The retinal image motion in these distraction conditions was comparable except for its origin, allowing for a direct examination of the impact of perceptually suppressed sensory inputs on VSTM.

Our results demonstrated that the delay-period distractor induced an attractive bias in VSTM regardless of its origin. This finding was also confirmed through cross-condition classification, where a classifier trained on data from one condition could predict observers’ responses in the other condition. An additional experiment further confirmed that the attractive bias induced by smooth pursuit was not an artifact of eye movement itself. These results not only provide behavioral evidence that even perceptually suppressed sensory inputs induced by self-action can act as a distractor for VSTM, but also suggest that a shared neural substrate between sensory and memory processing is independent of the ultimate perception.

## Results

The procedure for Experiment 1 is illustrated in Figure 1A (see Materials and Methods for details). In this experiment, human observers were asked to compare the speed of two motion stimuli presented sequentially (i.e., the sample and test motion), while ignoring task-irrelevant motion stimuli presented between them (i.e., the distractor motion). The distractor motion was presented either as retinal image motion with or without physical displacement (i.e., actual motion in world coordinate or apparent motion induced by eye movement). The motion stimuli consisted of a random-dot texture, with contrast modulated by a Gaussian window with a standard deviation (SD) of 0.96 (Figure 1B) (27, 28). The stimuli were presented in pairs, positioned 4.0 degrees (deg) above and below a fixation spot (a white Gaussian dot with SD of 0.15 deg). Observers were instructed not to gaze at the motion stimuli but at the fixation spot.

**Figure 1.**
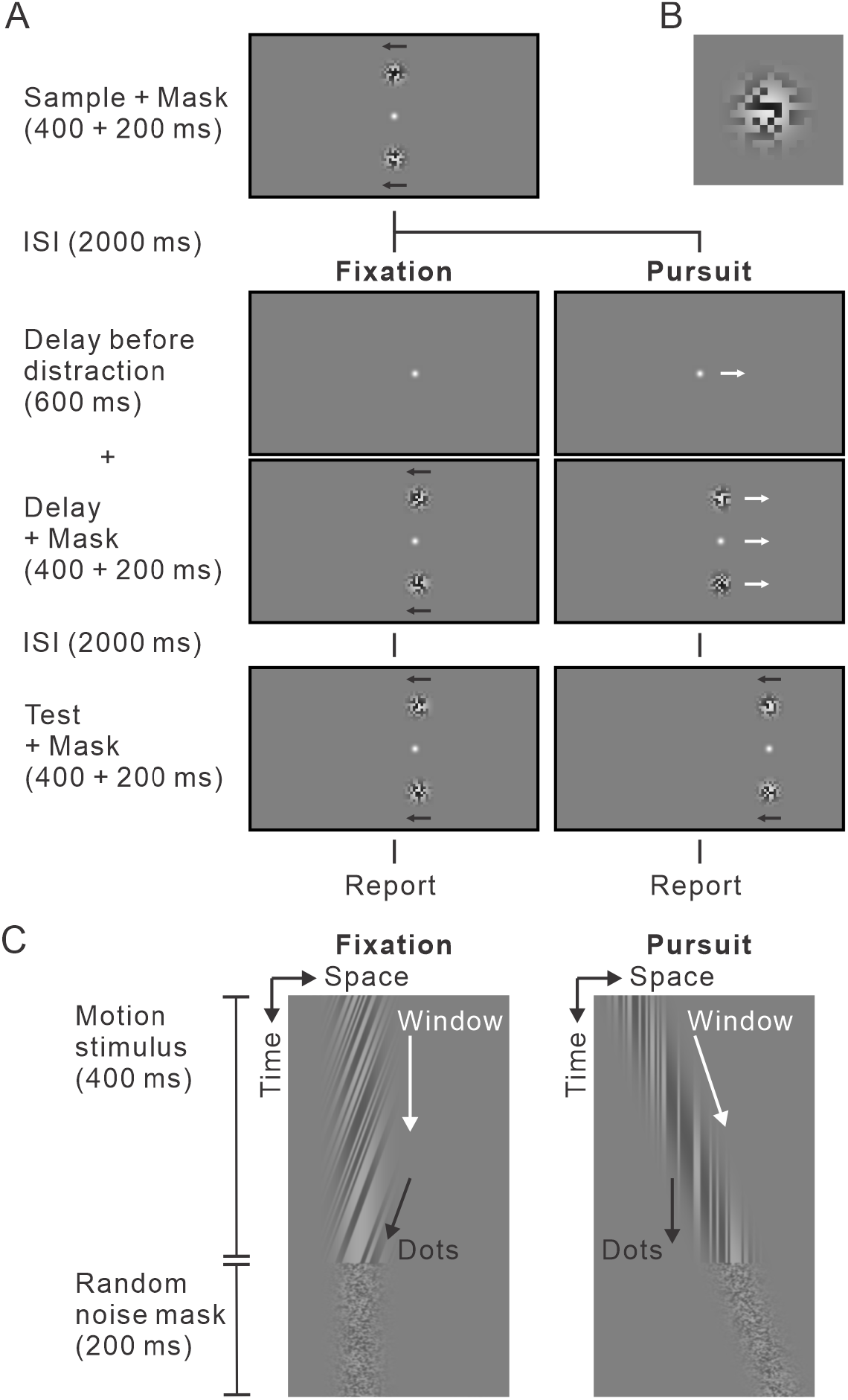
Visual short-term memory task with delay-period sensory distractor. (A) Trial sequence and conditions. The visual motion stimulus consisted of a random dots texture, with contrast modulated by a Gaussian window (see Figure 1B). Arrows depict the motion of the random-dot texture (black arrows) and the Gaussian window (white arrows). Observers were asked to compare the speed of sample and test motion stimuli. (B) Visual motion stimulus. The random-dot texture consisted of 50% black dots (luminance: 0.1 cd/m^2^) and 50% white dots (luminance: 70.0 cd/m^2^), each 4 × 4 pixels. (C) Spatiotemporal diagrams of motion stimuli during the delay period. In the fixation condition (left), the random-dot texture moved within the stationary Gaussian window, which is the same when presenting the sample and test motion (Figure 1A). In the pursuit condition (right), the fixation spot moved in the opposite direction to the sample motion, while the Gaussian window moved in the same direction over the stationary random-dot texture.

Both the sample and test motion were presented as the random-dot texture moving within the stationary Gaussian window, whereas the distractor was presented in one of the following two conditions. In the fixation condition, the distractor motion was presented in the same manner as the sample and test motion (i.e., motion in world coordinates, Figure 1C left). In the pursuit condition, the fixation spot moved in the opposite direction to the sample motion, and simultaneously the Gaussian window moved in the same direction over the stationary random-dot texture (Figure 1C right). This configuration ensured that in the pursuit condition, continuous ocular tracking of the fixation spot induced retinal image motion identical to the fixation condition, but without any motion in world coordinates. To minimize the influence of each motion stimulus on the perception of subsequent stimuli (i.e., motion aftereffect) (29), random noise masks were introduced at the end of each motion stimulus (Figure 1C) (30).

### “Retinal image motion distractor induces an attractive bias on VSTM for motion, even when generated by smooth pursuit”

We validated the distractor-induced bias in VSTM using generalized linear mixed models (GLMMs) applied to 3,353 trials from 8 observers. Three models with different combinations of fixed effects were designed to predict whether the test motion was perceived as faster than the sample motion (T^+^ response). Model 1 included the eye condition (the fixation or pursuit conditions), conditioned distractor speed (Δdist-speed: -2, ±0, or +2 deg/s relative to the sample speed), test speed (Δtest-speed: -2, -1, ±0, +1, or +2 deg/s relative to the sample speed), and their interactions as fixed effects. While this model reflected our experimental design, it might not fully capture the true effect, as the actual speed of retinal image motion depended on the speed of observers’ eye movement, particularly in the pursuit condition. Indeed, the pursuit gain varied across trials, with an average value of 0.88 (Figure S1A). In response, Model 2 included the eye condition, actual Δdist-speed (adjusted by eye speed [deg/s]), Δtest-speed, and their interactions as fixed effects. Model 3 included the actual Δdist-speed, Δtest-speed, and their interaction as fixed effects, hypothesizing that the distractor motion biases VSTM regardless of whether it is actual motion in world coordinates or apparent motion induced by smooth pursuit. We then compared 20 models with different combinations of random effects based on these three models (Figure S1B), ultimately selecting Model 3, which included random effects of actual Δdist-speed, Δtest-speed, and their interaction, based on the Akaike information criterion (AIC). The selected model revealed significant fixed effects of actual Δdist-speed (estimate: -0.16 [95% CI: -0.20, -0.12], *t*_3349_ = -8.47, *p* = 3.61 × 10^−17^) and Δtest-speed (estimate: 1.05 [95% CI: 0.96, 1.15], *t*_3349_ = 21.57, *p* = 9.93 × 10^−97^) with an *r*^2^ value of 0.92 (Figure 2A). The negative estimate of actual Δdist-speed indicates the attractive bias of VSTM induced by the distractor (6), corresponding to a systematic shift of the psychometric curve as a function of Δtest-speed (Figure 2B). Figure 2C summarizes the attractive bias induced by actual Δdist-speed, indexed by the point of subjective equality (PSE) computed by the psychometric curve. Overall, the PSE in the pursuit condition shifted in the negative direction, which could be explained by a decrease in the actual Δdist-speed due to the pursuit gain being less than 1.0.

**Figure 2.**
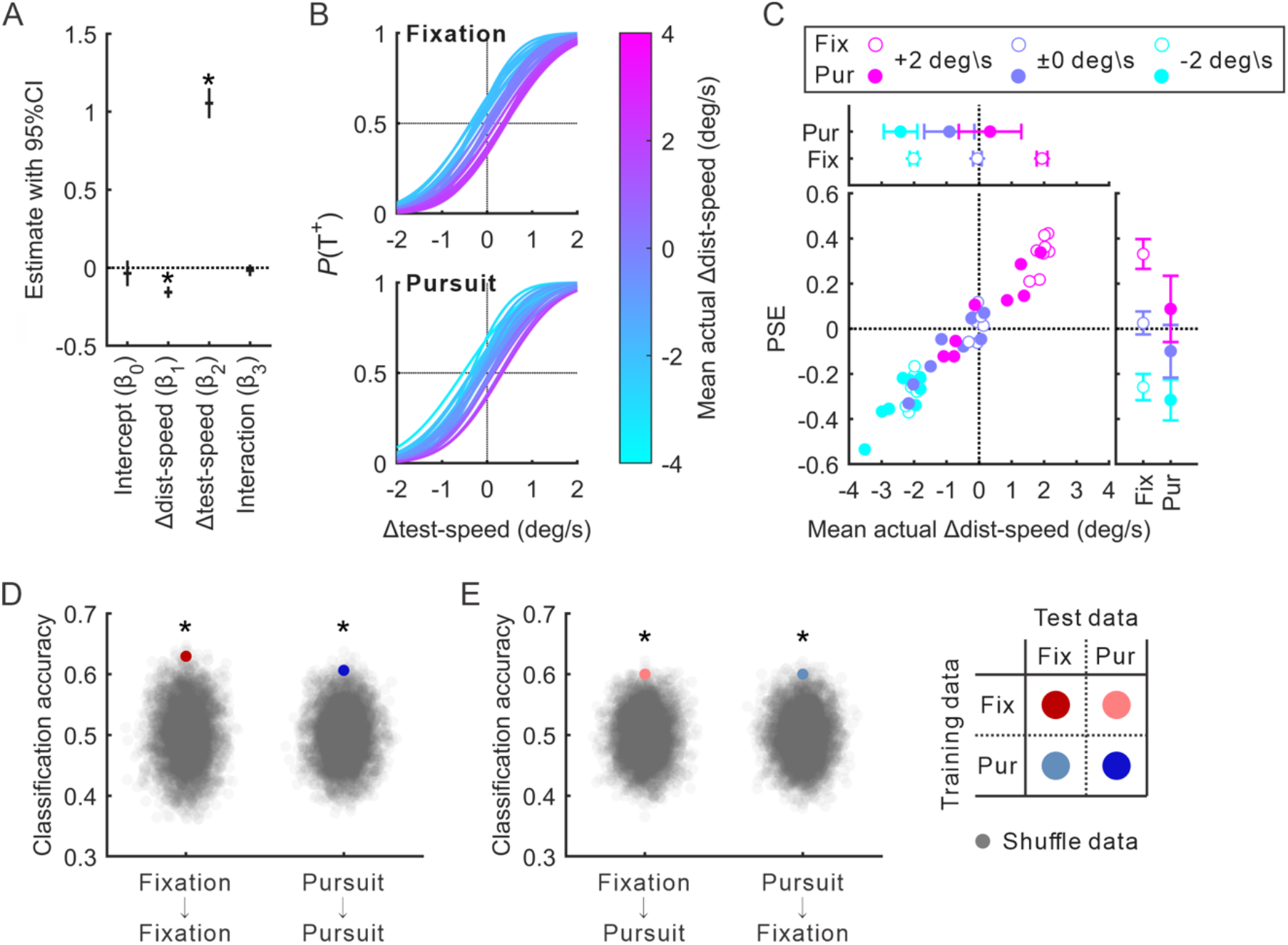
Summary of Experiment 1. (A) Fixed effect estimates and 95% CIs (vertical bars) for the selected GLMM (3,353 trials from 8 observers). The GLMM predicted whether the test motion was perceived as faster than the sample motion (T^+^ response) [response ∼ actual Δdist-speed * Δtest-speed + (1 + actual Δdist-speed * Δtest-speed | observer)]. Asterisks denote statistical significance (*p* < 0.05). actual Δdist-speed: distractor speed relative to sample speed, adjusted by eye speed (deg/s). Δtest-speed: test speed relative to sample speed (deg/s). (B) Psychometric curves as a function of Δtest-speed for the fixation (upper panel) and pursuit (lower panel) conditions. The mean actual Δdist-speed for each condition combination was calculated for each observer and then inputted into the GLMM to generate 48 psychometric curves (2 eye conditions × 3 Δdist-speed conditions × 8 observes). (C) Relationship between the PSE and mean actual Δdist-speed. The scatter plot displays a total of 48 data points corresponding to the psychometric curves in Figure 2B. The top panel shows the mean ± 95% CI of mean actual Δdist-speed for the fixation (Fix) and pursuit (Pur) conditions. The right panel shows the mean ± 95% CI of PSE for both conditions. (D) Performance of within-condition classification. The classifier was trained and tested within each condition. The colored dot represents the median performance using true data, whereas the gray dots represent the distribution of performance from label-shuffled data (n = 5,000). Asterisks denote statistical significance (permutation test. *p* < 0.05). (E) Performance of cross-condition classification, plotted in the same format as Figure 2D. The classifier was trained on one condition and tested on the other condition.

To quantify the extent to which the observer’s response was influenced by the distractor and to further confirm the independence of the distraction effect from its origin, we used logistic regression to classify the observers’ responses based on the actual Δdist-speed. We focused on no-difference trials (i.e., trials where Δtest-speed = 0), where observers were expected to make random guesses. We trained a classifier on the dataset from one condition and tested its performance on both the same condition (within-condition classification) and the other condition’s dataset (cross-condition classification). If, as suggested by the GLMM, retinal image motion induces the attractive bias in VSTM regardless of its origin, the classifier should be able to predict the observers’ responses in trials from the other condition that was not used for training. The performance of the within-condition classification was 0.63 for the fixation condition and 0.61 for the pursuit condition (Figure 2D). Significance was confirmed by permutation tests using an empirical null distribution based on 5,000 iterations of classification on label-shuffled data (fixation: *p* = 1.80 × 10^−3^; pursuit: *p* = 3.00 × 10^−3^). These modest but significant effects correspond to previous studies on the attractive bias in VSTM for other visual features (7–9). Similar results were observed in the cross-condition classification, where the fixation-condition-trained classifier performed at 0.60 on the pursuit condition’s dataset (permutation test, *p* = 1.60 × 10^−3^, Figure 2E left), and at 0.60 for the reverse case (permutation test, *p* = 3.60 × 10^−3^, Figure 2E right). The significant performance in the cross-condition classification indicates that the observers’ responses were, at least in part, biased by the actual Δdist-speed in the same manner across both conditions. These results further emphasize the attractive bias in VSTM induced by the delay-period retinal image motion, regardless of its origin.

### “Eye movement alone does not function as a distractor”

In Experiment 1, since the speed of distractor motion in the pursuit condition corresponded to the speed of smooth pursuit, we could not rule out the possibility that eye movement itself functions as a distractor for VSTM. Some studies have mentioned that eye movement or its control during the delay period can distract VSTM for locations (31, 32). Furthermore, a subpopulation of MST neurons exhibits firing activity related to smooth pursuit without substantial visual motion (33–35). Therefore, Experiment 2 was designed to test the effect of eye movement on the VSTM bias while removing retinal image motion. The experimental procedure in Experiment 2 was largely identical to the pursuit condition in Experiment 1, except that the random-dot texture and the Gaussian window moved coherently in the same direction, resulting in no retinal image motion during smooth pursuit (Figure 3A). If the attractive bias observed in the pursuit condition of Experiment 1 was driven by eye movement itself rather than retinal image motion, the distraction effect should persist in Experiment 2.

**Figure 3.**
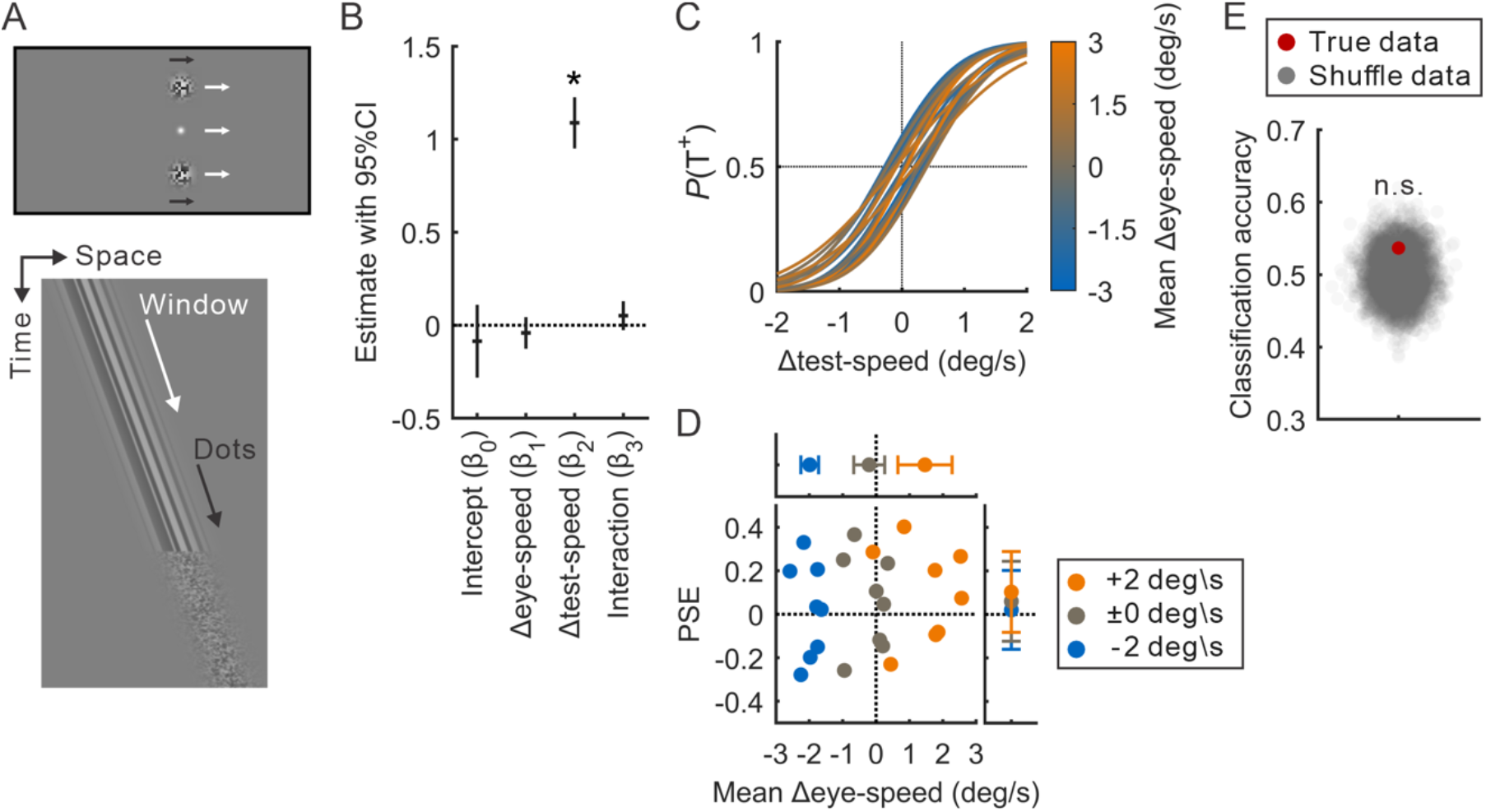
Summary of Experiment 2. (A) Visual motion stimulus during the delay period (upper panel) and spatiotemporal diagram of the motion (lower panel). The experimental procedure in Experiment 2 was largely identical to the pursuit condition in Experiment 1. When observers tracked the moving fixation spot with their gaze, the random-dot texture and the Gaussian window moved coherently in the same direction. (B) Fixed effect estimates and 95% Cis for the selected GLMM (1,706 trials from 8 observers). The GLMM predicted whether the test motion was perceived as faster than the sample motion (T^+^ response) [response ∼ Δeye-speed * Δtest-speed + (1 + Δeye-speed * Δtest-speed | observer)]. An asterisk denotes statistical significance (*p* < 0.05). Δeye-speed: eye speed relative to sample speed (deg/s). Δtest-speed: test speed relative to sample speed (deg/s). (C) Psychometric curves as a function of Δtest-speed. The mean Δeye-speed for each condition combination was calculated for each observer and then inputted into the GLMM to generate 24 psychometric curves (3 Δeye-speed conditions × 8 observes). (D) Relationship between the PSE and mean Δeye-speed, plotted in the same format as Figure 2C. (E) Performance of classification, plotted in the same format as Figure 2D.

Here, three models with different combinations of fixed effects were designed to predict T^+^ response (1706 trials from 8 observers). The first model was comparable to Model 3 in Experiment 1, which included the actual Δdist-speed, Δtest-speed, and their interaction as fixed effects. Note that the actual Δdist-speed in Experiment 2 was determined sorely by pursuit gain (Figure S2A) and was totally smaller than that in Experiment 1. Model 2 included the Δeye-speed (difference between the sample motion and eye speed during smooth pursuit [deg/s]), Δtest-speed, and their interaction as fixed effects, which did not include retinal image motion of the distractor directly as fixed effects. Finally, Model 3 included both variables regarding retinal and eye information, that is, the actual Δdist-speed, Δeye-speed, Δtest-speed, and their interactions as fixed effects. Based on the AIC value, Model 2, which included random effects of the Δeye-speed, Δtest-speed, and their interaction, was selected from 16 models with different combinations of random effects (Figure S2B). This model revealed a significant fixed effect of Δtest-speed (estimate: 1.07 [95% CI: 0.94, 1.20], *t*_1702_ = 15.75, *p* = 2.79 × 10^−52^) but no significance in Δeye-speed (estimate: -0.03 [95% CI: -0.07, 0.02], *t*_1702_ = -1.02, *p* = 0.31), with an *r*^2^ value at 0.92 (Figure 3B). Indeed, Δeye-speed no longer explained the shift of the psychometric curve (Figure 3C and 3D).

We also computed a classifier that predicts observers’ responses in no-difference trials. The performance of the classifier trained using Δeye-speed as a predictor did not significantly differ from chance level, as determined by a permutation test (*p* = 0.11. Figure 3E). These results suggest that eye movement alone does not function as a distractor to bias VSTM and confirm that the findings in Experiment 1, namely the attractive bias were induced by retinal image motion during smooth pursuit.

## Discussion

Given previous studies that reported no effect of unconscious sensory distractors on VSTM (24, 25), our results, which show the distraction effect of perceptually suppressed sensory inputs, are somewhat surprising. The discrepancy between our findings and previous studies may be attributed to several factors. First, sensory inputs from unconscious visual stimuli in previous studies might be insufficient to induce a VSTM bias (24, 25). We addressed this potential issue by aligning key visual stimulus properties, including presentation duration, between the conditions. Moreover, previous studies on the effect of unconscious sensory distractors on VSTM has focused primarily on VSTM for orientation, which is mainly processed via the ventral stream, in contrast to the dorsal stream for visual motion processing (36). Additionally, the mechanisms underlying perceptual suppression differ: for example, binocular rivalry is associated with reduced neural activity in the ventral stream regarding suppressed information (26), whereas perceptual invariance during smooth pursuit is achieved by integrating retinal and extraretinal signals after encoding (15, 16). Furthermore, the impact of unconscious distractors has been found to be inconsistent, likely due to variations in experimental procedures. For example, subliminal distractors with brief presentation durations influence VSTM performance when presented during the delay period (37), but not before the sample period (24). Taken together, our findings provide behavioral evidence that retinal image motion during the delay period can act as a sensory distractor to VSTM for motion without conscious perception, though further studies are needed to explore its applicability to VSTM for other visual features.

Perceptual compensation during smooth pursuit is known to be incomplete (15, 16), which raises the possibility that the distraction effect observed in the pursuit condition of Experiment 1 could be due to residual motion signals that yield a visual motion illusion (38). One potential illusion involved in our task is the Filehne illusion (39), where stationary background objects are perceived to move in the opposite direction during smooth pursuit. The Filehne illusion depends on both eye speed during smooth pursuit and the duration of background object presentation (40, 41), and our experimental procedure might not fully rule out this illusion. If residual motion signals and/or related perceptual illusions were responsible for our results, we would expect the impact to be considerably smaller than the expected retinal image motion. However, the observed impact was comparable to that in the fixation condition, as confirmed by the GLMM and cross-condition classification. We therefore conclude that residual motion signals and/or related perceptual illusions are unlikely to account for our results.

Our findings also suggest a dissociation between neural substrates involved in VSTM and visual perception. Here we discuss the underlying mechanisms of the sensory-memory interference independent of ultimate perception. Note that we assume that this interference originates in higher brain regions, such as the PFC and the posterior parietal cortex (PPC), rather than in early visual areas (42). While the role of early visual areas in memory maintenance remains debated, recent studies have shown that these areas encode mnemonic and sensory information through distinct neural populations or formats (22, 43, 44) and suggested that the mnemonic decodability of BOLD signals in early visual areas (45–47) reflects a top-down mechanism by memory signals from the PFC (22, 48). The coexistence of mnemonic and sensory representations can be achieved by layer-specific inputs (observed in V1 (49, 50)), different encoding formats, and subthreshold potentiation (22, 51). Importantly, the top-down memory signal could serve as a template for comparing with bottom-up sensory inputs, shaping a local comparison circuit within early visual areas (11, 52). This idea is supported by observations that distractor-induced behavioral biases are predicted by the mnemonic representation in early sensory areas, likely driven by the top-down signal (11, 52). The mechanism appears ideal for typical VSTM tasks, such as change detection and discrimination, and is also applicable to VSTM for motion (14, 53). Based on these findings, we consider it plausible to focus on the PFC and PPC which are involved in distraction resistance in VSTM (54–56).

The representation of visual motion is transformed from retinal to world coordinates along the hierarchy of the dorsal stream. For example, while MT neurons predominantly encode motion in retinal coordinates, a subset of MST neurons exhibits motion signals in world coordinates, which are more directly related to perceptual experience (18–20). The dorsal stream projects to the PPC (36), which is also known to encode VSTM with distraction resistance (54–56). If mnemonic representations in the PPC rely on perceptually compensated sensory signals inherited from the dorsal stream, and this interaction causes a VSTM bias, the bias should be correlated with ultimate perception rather than original retinal signals. However, this prediction contradicts our behavioral results, where the bias was linked to not perception, but to sensory signals (i.e., retinal image motion). This discrepancy may be explained by the mechanism for perceptual invariance during smooth pursuit is supported by distributed networks, rather than the dorsal stream alone (16, 57). Even within the MST, which has recently been implicated in the distraction resistance of VSTM for motion through a computational model involving recurrent neural networks (58), the majority of neurons still encode motion in retinal coordinates rather than world coordinates (59). The large number of neurons related to perceived motion are also found in the visual posterior sylvian (VPS) area of monkeys, where a similar proportion of neurons encode perception and retinal image motion (60). VPS, located in the caudal lateral sulcus below the classical dorsal stream, receives inputs from the MST (61), and human studies have found that a putative human homologues of VPS exhibits the perception-related activity during smooth pursuit (15, 62). Remarkably, a report describes a patient with bilateral lesions in parts of the parieto-occipital cortex, possibly including the putative human homologue of VPS, who perceived smooth pursuit-induced retinal image motion as motion in the world, though V1 and most probably at least parts of the MT/MST were available (63). Outside of VPS, activities related to perceptual invariance during smooth pursuit have been observed in the cerebrocerebellar circuit and the parietoinsular vestibular cortex (PIVC) (64). In other words, the visual motion information that reaches the PPC may be close to the original retinal signals for which perceptual compensation is insufficient. Further, the influence could be transmitted to the PFC via the PFC-PPC circuit with bidirectional connections (42, 65). Taken together, we hypothesize that the PFC-PPC circuit, which converts visual motion signals into VSTM, relies on sensory signals before compensation for eye movement, resulting in that even perceptually suppressed sensory inputs interfere with VSTM. This hypothesis may be tested in future neurophysiological studies that determine whether mnemonic representations in the PFC-PPC circuit reflect sensory or perceptual information.

Finally, does the dissociation between VSTM and visual perception confer any advantage in neural computation? One possibility is that the local circuit in early visual areas which compares bottom-up sensory signals with top-down memory signals may prefer encoding memorized items in the same format as the original sensory inputs. If this is the case, sensory distractors that are not perceived may not influence the memory when visual stimuli are memorized as semantic information (66). Another possibility is that representing VSTM as sensory rather than perceptual information could be advantageous in more complex cognitive tasks including manipulating memorized information, compared to simpler tasks like comparing pairs of sequentially presented stimuli, as done in this study. This could include mental rotation or transformations of memorized items and reference frame transformation from retinal to other spatial coordinates (67).

In summary, our results demonstrate that retinal image motion induced by smooth pursuit can distract VSTM for motion, even though the motion signal is perceptually suppressed. This suggests that the sensory-memory interference is independent of the ultimate motion perception. This dissociation between VSTM and perception likely stems from the independence of systems for perceptual invariance during smooth pursuit and encoding VSTM based on sensory inputs. Although perception is believed as a process of active inference about the external world based on sensory evidence (68), most VSTM studies have not clearly distinguished between perceptual experience and sensory inputs (the potential importance is mentioned in the review by Roussy et al. (23)). Our findings provide behavioral evidence that sensory inputs can distract VSTM without conscious perception, while also suggest that the VSTM system shares neural substrates with sensory processing, but not with perception. At the same time, our findings also emphasize the need to distinguish between sensory signals and perceptual signals in VSTM research.

## Materials and Methods

### Observers

Eight observers (8 men. 24.8 ± 2.1 years, mean ± SD) participated in Experiment 1, and eight (7 men and 1 woman. 26.0 ± 2.7 years, mean ± SD) observers participated in Experiment 2. They reported having normal or corrected-to-normal vision and no known visuomotor deficits. Sample sizes were not statistically predetermined but were comparable to those in previous studies on the distraction effect of unconscious sensory distractors (24, 25). They provided written informed consent in accordance with the Declaration of Helsinki and were informed of their right to withdraw from the study at any time without penalty. The study protocol was approved by the Ethics Committee of the Department of Cognitive and Psychological Sciences at Nagoya University (No. 240801-C-04-1).

### General settings

Observers were seated 57.0 cm from an LCD monitor (AW2524HF; Dell Technologies Inc., Texas, USA; size: 24.5 inches, resolution: 1920 × 1080 pixels, refresh rate: 120 Hz) with their heads stabilized by a chin rest and forehead restraint. The monitor was positioned with its height adjusted so that the observer’s eye level was aligned with a fixation spot during the task.

Eye movement of the right eye was recorded using a video-based eye-tracking system. Eye position signals were detected via reflections of infrared light on the cornea, and the pupil’s image was captured by an infrared camera (GS3-U3-41C6NIR; FLIR Systems Inc., Oregon, USA) (69). The system digitized eye position signals at 1 kHz with 12-bit precision using an A/D converter (TUSB-1612ADSM-S2Z; Turtle Industry Co., Ltd., Ibaraki, Japan). Prior to the task, eye position signals were calibrated by having observers fixate on target spots (diameter: 0.3 deg, luminance: 70.0 cd/m^2^) at known horizontal and vertical eccentricities under binocular viewing conditions. All stimuli during calibration and the main task were presented on a uniform gray background (luminance: 17.0 cd/m^2^).

The visual motion stimuli consisted of a random-dot texture, with contrast modulated by a Gaussian window (SD: 0.96) (Figure 1B) (27, 28). This corresponded to the size at which the random-dot texture of about 2.4 deg can be seen at a contrast of 20% or more. The random-dot texture consisted of 50% black dots (luminance: 0.1 cd/m^2^) and 50% white dots (luminance: 70.0 cd/m^2^), each 4 × 4 pixels. All visual stimuli and experimental routine were programmed using Psychophysics Toolbox extensions (70–72) in MATLAB (MathWorks, Massachusetts, USA).

### Tasks and Procedure

#### Experiment 1

Each trial began with a fixation spot (a white Gaussian dot with SD of 0.15 deg) displayed in the center of the monitor and a text instruction (“Press any key to continue”) above it. Observers were instructed to maintain their gaze on the fixation spot throughout the trial.

When observers pressed any key on a handheld numeric keypad, the text disappeared, followed by a 1.2-1.5 s fixation period. Afterward, the motion stimuli were presented in pairs, positioned 4.0 deg above and below the fixation spot. Each random-dot texture moved either to the right or left within the stationary Gaussian window for 0.4 s, with speed randomly selected from 6.0, 7.0, and 8.0 deg/s (sample period, Figure 1A). The motion direction reversed each trial. Immediately after the stimulus offset, random noise masks, in which each dot was randomly replaced every 0.01 s, were presented for 0.2 s. This was introduced to minimize the motion aftereffect and was applied after all motion stimuli in this study. Following the mask, an additional 2.0 s fixation period was provided. Between 1.0 and 1.7 s into the fixation period, a text instruction was displayed above the fixation spot (i.e., “Fixation” or “Pursuit”), informing observers of the required eye movement for the upcoming motion stimulus (distractor period, Figure 1A). In the fixation condition, the fixation spot disappeared and represented at an offset location 1.5-2.0 deg in the opposite direction of the sample motion. Observers were instructed to refocus on the new fixation spot, mimicking a catch-up saccade that typically occurs at the onset of ocular tracking in the pursuit condition. This was also aimed to prevent spatial overlap of the sample and distractor motion in screen coordinates. After 0.6 s since the fixation spot reappeared, paired motion stimuli were presented, similar to the sample motion, but with speed selected from -2.0, ±0.0, and +2.0 deg/s relative to the sample speed (Figure 1C left). In the pursuit condition, observers were instructed to track the fixation spot that moved in the opposite direction of the sample motion, at a constant speed selected from -2.0, ±0.0, and +2.0 deg/s relative to the sample speed. After 0.6 s of the fixation translation onset, paired motion stimuli were presented, where the Gaussian window moved at the same speed and direction as the fixation spot over the stationary random-dot texture, resulting in retinal image motion without physical displacement of the random-dot texture (Figure 1C right). The fixation spot stopped after 0.6 s (0.4 s of motion stimulus plus 0.2 s of mask). In both conditions, after an additional 2.0 s fixation period, paired motion stimuli were presented again, similar to the sample motion, with speed selected from -2.0, -1.0, ±0.0, +1.0, and +2.0 deg/s relative to the sample speed (test period, Figure 1A). After 1.0 s of blank, observers reported whether the test motion was perceived as faster (T^+^ response) or not (T^-^ response) compared to the sample motion using the keypad (pressing the “6” key for T^+^ and the “4” key for T^-^). No performance feedback was provided to observers.

Each observer completed 450 trials (2 eye conditions × 3 distractor speeds × 5 test speeds × 15 repetitions), interleaved and divided into ten blocks. For familiarization, observers practiced the task for one or two blocks prior to the main task.

#### Experiment 2

The experimental procedure was largely identical to that of the pursuit condition in Experiment 1, except that the random-dot texture and the Gaussian window moved coherently in the same direction, resulting in no retinal image motion during smooth pursuit (Figure 3A). The speed of the moving fixation spot during the distractor period was selected from -2.0, ±0.0, and +2.0 deg/s relative to the sample speed, similar to the procedure in Experiment 1. Each observer completed 225 trials (3 distractor speeds × 5 test speeds × 15 repetitions), interleaved and divided into ten blocks. For familiarization, observers practiced the task for one or two blocks prior to the main task.

### Quantification and Statistical analysis

The method for analyzing eye movement data was based on previous research (73, 74). Eye position data were filtered with a second-order Butterworth low-pass filter with a 15 Hz passband. Eye velocity and acceleration were calculated using digital differentiation of the position data with the central difference algorithm and then filtered with a second-order Butterworth low-pass filter with a 30 Hz passband. Saccades were identified based on criteria of acceleration exceeding 1000 deg/s^2^ and velocity exceeding 30 deg/s. Eye speed was defined as the mean eye velocity during the distractor period. Pursuit gain was the ratio of eye speed to the speed of the fixation spot translation. The sign for all motion (including stimuli and eye movement) was defined as positive to the right.

Motion speed variables were defined relative to the speed of the sample motion. Δdist-speed represented the speed of the distractor motion relative to the sample motion (i.e., -2.0, ±0.0, and +2.0 deg/s), and Δtest-speed represented the speed of test motion relative to the sample motion (i.e., -2.0, -1.0, ±0.0, +1.0, and +2.0 deg/s). Additionally, we calculated the actual speed of distractor by subtracting eye speed from the distractor speed in world coordinates (i.e., provided distractor speed minus eye speed in the fixation condition, and zero minus eye speed in the pursuit condition). The actual Δdist-speed was then defined as the actual speed of distractor relative to the sample speed.

#### Experiment 1

Trials were excluded if observers blinked during any motion stimuli, made gaze shifts of 1.0 deg or more from the fixation spot during the sample and test periods (in both conditions) or during the distractor period (in the fixation condition), made saccades during the distractor period in the pursuit condition, or had a pursuit gain less than 0.5 or greater than 1.5 in the pursuit condition. A total of 3353 out of 3600 trials (93.1%) were included in the analysis.

To assess the impact of delay-period perceptual distractors on VSTM performance, we used generalized linear mixed models (GLMMs) with a probit link function to predict T^+^ responses. We constructed 20 models combining the following three models with random effects (all models included random intercept). Model 1 included the eye condition, Δdist-speed, Δtest-speed, and their interactions as fixed effects. Model 2 included the eye condition, actual Δdist-speed, Δtest-speed, and their interactions as fixed effects. Model 3 included the actual Δdist-speed, Δtest-speed, and their interaction as fixed effects. Model selection was based on the Akaike information criterion (AIC), and Model 3 with the actual Δdist-speed, Δtest-speed, and their interaction as random effects was selected as the best model. Note that the result of the model selection was the same when based on the Bayesian information criterion (BIC) (data not shown). Estimates of fixed effects of this model are shown in Figure 2A, and statistical significance denoted as a single asterisk (*p* < 0.05).

Using the selected model, we generated psychometric curves as a function of Δtest-speed.

The mean actual Δdist-speed for each combination of eye conditions and Δdist-speed conditions was calculated for each observer and input into the model to generate 48 psychometric curves (2 eye conditions × 3 Δdist-speed conditions × 8 observes; Figure 2B). The point of subjective equality (PSE) for each curve was calculated as an index of the distractor-induced bias in VSTM and plotted against the mean actual Δdist-speed (Figure 2C). Dots and error bars outside the scatter plot in Figure 2C show the mean and 95% confidence interval (CI) for the mean actual Δdist-speed (in x axis) and PSE (in y axis) for each combination across 8 observers.

To quantify the influence of the distractor on observers’ responses and further confirm the independence of the distraction effect from its origin, we performed logistic regression on no-difference trials (where Δtest-speed = 0) to classify observers’ responses based on actual Δdist-speed. We trained a classifier on the dataset from one condition and tested its performance on both the same condition (within-condition classification) and the other condition’s dataset (cross-condition classification). The fixation condition included 180 trials for T^+^ responses and 174 trials for T^-^ responses, and the pursuit condition included 160 trials for T^+^ responses and 166 trials for T^-^ responses. To balance the number of trials across all labels, we randomly sampled 160 trials from each label. Using these trials, we conducted a stratified 10-fold cross-validation procedure in each condition, where a classifier was trained on 9 out of 10 splits (i.e., 144 trials for each of the T^+^ and T^-^ responses) and tested on both the remaining one split and one split of the other condition (i.e., 16 trials for each response in each condition). This procedure was also applied to classifiers trained with shuffled data, where class labels were randomly permutated. This process was repeated 10 times with different splits for test dataset, and the accuracy rate between classifiers’ outputs and observers’ responses was used to evaluate the performance of each iteration. The above series of steps was repeated 5000 times with different sample data, and the overall performance of each classifier for each condition was defined as the median performance across all iterations. Significance of the overall performance was determined using a permutation test with the distribution of corresponding shuffled data (*p* < 0.05) (75), as indicated by a single asterisk in Figure 2D and 2E.

#### Experiment 2

Invalid trials were identified and excluded based on the same criteria in th pursuit condition of Experiment 1, resulting in a total of 1706 out of 1800 trials (94.8%) were included in the analysis. Although the direction of eye movement was always opposite to that of the sample motion, to maintain consistency with Experiment 1, Δeye-speed was calculated using the absolute values of the eye and sample speeds.

To assess the impact of eye speed during the distractor period on VSTM performance, GLMMs were used to predict T^+^ responses. We constructed 16 models combining the following three models with random effects. Model 1 included the actual Δdist-speed, Δtest-speed, and their interaction as fixed effects, identical to Model 3 in Experiment 1. Model 2 included the Δeye-speed, Δtest-speed, and their interaction as fixed effects. Model 3 included the actual Δdist-speed, Δeye-speed, Δtest-speed, and their interactions as fixed effects. Model selection based on the AIC identified Model 2 with the Δeye-speed, Δtest-speed, and their interaction as random effects as the best model. Estimates of fixed effects of this model are shown in Figure 3B, and statistical significance denoted as a single asterisk (*p* < 0.05). Similar to Experiment 1, psychometric curves as a function of Δtest-speed were generated by inputting the mean Δeye-speed for each of three Δdist-speed conditions (24 curves from 8 observers; Figure 3C), and the PSE was plotted in Figure 3D. Classification analysis was conducted using the similar procedure of within-condition classification in Experiment 1, but in this case, the classifier predicted observers’ responses based on Δeye-speed (Figure 3E).

## Data and Code Availability

All data and original codes have been deposited at Zenodo (doi: 10.5281/zenodo.15123600) and are publicly available as of the date of publication. Any additional information required to reanalyze the data reported in this paper is available from the corresponding author (T.M.) upon request.

## Acknowledgments

This study was supported by JSPS KAKENHI (grant no.: 22KJ1787 and 23K16671), JST CREST (grant no.: JPMJCR22P5), and Open Access Acceleration Project.

## References

1. A. Baddeley, Working memory. Curr. Biol. 20, 136–140 (2010).

2. D. Soto, J. Hodsoll, P. Rotshtein, G. W. Humphreys, Automatic guidance of attention from working memory. Trends Cogn. Sci. 12, 342–348 (2008).

3. F. van Ede, Visual working memory and action: Functional links and bi-directional influences. Vis. cogn. 28, 401–413 (2020).

4. F. McNab, R. J. Dolan, Dissociating distractor-filtering at encoding and during maintenance. J. Exp. Psychol. Hum. Percept. Perform. 40, 960–967 (2014).

5. H. R. Liesefeld, A. M. Liesefeld, P. Sauseng, S. N. Jacob, H. J. Müller, How visual working memory handles distraction: cognitive mechanisms and electrophysiological correlates. Vis. cogn. 28, 372–387 (2020).

6. E. S. Lorenc, R. Mallett, J. A. Lewis-Peacock, Distraction in visual working memory: Resistance is not futile. Trends Cogn. Sci. 25, 228–239 (2021).

7. S. Magnussen, M. W. Greenlee, R. Asplund, S. Dyrnes, Stimulus-specific mechanisms of visual short-term memory. Vision Res. 31, 1213–1219 (1991).

8. D. J. McKeefry, M. P. Burton, C. Vakrou, Speed selectivity in visual short term memory for motion. Vision Res. 47, 2418–2425 (2007).

9. J. H. Yoon, C. E. Curtis, M. D’Esposito, Differential effects of distraction during working memory on delay-period activity in the prefrontal cortex and the visual association cortex. Neuroimage 29, 1117–1126 (2006).

10. T. Pasternak, M. W. Greenlee, Working memory in primate sensory systems. Nat. Rev. Neurosci. 6, 97–107 (2005).

11. G. E. Hallenbeck, T. C. Sprague, M. Rahmati, K. K. Sreenivasan, C. E. Curtis, Working memory representations in visual cortex mediate distraction effects. Nat. Commun. 12, 4714 (2021).

12. S. A. Harrison, F. Tong, Decoding reveals the contents of visual working memory in early visual areas. Nature 458, 632–635 (2009).

13. D. Mendoza-Halliday, J. C. Martinez-Trujillo, Neuronal population coding of perceived and memorized visual features in the lateral prefrontal cortex. Nat. Commun. 8, 15471 (2017).

14. D. Zaksas, T. Pasternak, Directional signals in the prefrontal cortex and in area MT during a working memory for visual motion task. J. Neurosci. 26, 11726–11742 (2006).

15. P. Thier, T. Haarmeier, S. Chakraborty, A. Lindner, A. Tikhonov, Cortical substrates of perceptual stability during eye movements. Neuroimage 14, 33–39 (2001).

16. M. Furman, M. Gur, And yet it moves: Perceptual illusions and neural mechanisms of pursuit compensation during smooth pursuit eye movements. Neurosci. Biobehav. Rev. 36, 143–151 (2012).

17. R. T. Born, D. C. Bradley, Structure and function of visual area MT. Annu. Rev. Neurosci. 28, 157–189 (2005).

18. N. Inaba, S. Shinomoto, S. Yamane, A. Takemura, K. Kawano, MST neurons code for visual motion in space independent of pursuit eye movements. J. Neurophysiol. 97, 3473– 3483 (2007).

19. L. Chukoskie, J. A. Movshon, Modulation of visual signals in macaque MT and MST neurons during pursuit eye movement. J. Neurophysiol. 102, 3225–3233 (2009).

20. U. J. Ilg, S. Schumann, P. Thier, Posterior parietal cortex neurons encode target motion in world-centered coordinates. Neuron 43, 145–151 (2004).

21. S. Ono, The neuronal basis of on-line visual control in smooth pursuit eye movements. Vision Res. 110, 257–264 (2015).

22. D. Mendoza-Halliday, S. Torres, J. C. Martinez-Trujillo, Sharp emergence of feature-selective sustained activity along the dorsal visual pathway. Nat. Neurosci. 17, 1255–1262 (2014).

23. M. Roussy, D. Mendoza-Halliday, J. C. Martinez-Trujillo, Neural substrates of visual perception and working memory: Two sides of the same coin or two different coins? Front. Neural Circuits 15, 764177 (2021).

24. T. Wildegger, N. E. Myers, G. Humphreys, A. C. Nobre, Supraliminal but not subliminal distracters bias working memory recall. J. Exp. Psychol. Hum. Percept. Perform. 41, 826– 839 (2015).

25. R. L. Rademaker, I. M. Bloem, P. De Weerd, A. T. Sack, The impact of interference on short-term memory for visual orientation. J. Exp. Psychol. Hum. Percept. Perform. 41, 1650–1665 (2015).

26. F. Tong, M. Meng, R. Blake, Neural bases of binocular rivalry. Trends Cogn. Sci. 10, 502– 511 (2006).

27. M. J. Hawken, K. R. Gegenfurtner, Pursuit eye movements to second-order motion targets. J. Opt. Soc. Am. A 18, 2282–2296 (2001).

28. T. Miyamoto, K. Miura, T. Kizuka, S. Ono, Properties of smooth pursuit and visual motion reaction time to second-order motion stimuli. PLoS One 15, e0243430 (2020).

29. S. Anstis, F. A. J. Verstraten, G. Mather, The motion aftereffect. Trends Cogn. Sci. 2, 111–117 (1998).

30. N. Zokaei, S. Manohar, M. Husain, E. Feredoes, Causal evidence for a privileged working memory state in early visual cortex. J. Neurosci. 34, 158–162 (2014).

31. B. R. Postle, C. Idzikowski, S. Della Sala, R. H. Logie, A. D. Baddeley, The selective disruption of spatial working memory by eye movements. Q. J. Exp. Psychol. 59, 100–120 (2006).

32. D. G. Pearson, A. Sahraie, Oculomotor control and the maintenance of spatially and temporally distributed events in visuo-spatial working memory. Q. J. Exp. Psychol. A 56, 1089–1111 (2003).

33. U. J. Ilg, P. Thier, Visual tracking neurons in primate area MST are activated by smooth-pursuit eye movements of an “imaginary” target. J. Neurophysiol. 90, 1489–1502 (2003).

34. S. Ono, M. J. Mustari, Extraretinal signals in MSTd neurons related to volitional smooth pursuit. J. Neurophysiol. 96, 2819–2825 (2006).

35. S. Ono, M. J. Mustari, Role of MSTd extraretinal signals in smooth pursuit adaptation. Cereb. Cortex 22, 1139–47 (2012).

36. M. A. Goodale, A. D. Milner, Separate visual pathways for perception and action. Trends Neurosci. 15, 20–25 (1992).

37. J. Silvanto, D. Soto, Causal evidence for subliminal percept-to-memory interference in early visual cortex. Neuroimage 59, 840–845 (2012).

38. M. Spering, A. Montagnini, Do we track what we see? Common versus independent processing for motion perception and smooth pursuit eye movements: A review. Vision Res. 51, 836–852 (2011).

39. W. U. Filehne, Über das optische Wahrnehmen von Bewegungen. Zeitschrift für Sinnesphysiology 53, 134–145 (1922).

40. J. L. Souman, I. T. C. Hooge, A. H. Wertheim, Vertical object motion during horizontal ocular pursuit: compensation for eye movements increases with presentation duration. Vision Res. 45, 845–853 (2005).

41. A. H. Wertheim, Retinal and extraretinal information in movement perception: How to invert the Filehne illusion. Perception 16, 299–308 (1987).

42. Y. Xu, Reevaluating the sensory account of visual working memory storage. Trends Cogn. Sci. 21, 794–815 (2017).

43. J. Huang, et al., Neuronal representation of visual working memory content in the primate primary visual cortex. Sci. Adv. 10, eadk3953 (2024).

44. S. E. Favila, B. A. Kuhl, J. Winawer, Perception and memory have distinct spatial tuning properties in human visual cortex. Nat. Commun. 13, 5864 (2022).

45. A. C. Riggall, B. R. Postle, The relationship between working memory storage and elevated activity as measured with functional magnetic resonance imaging. J. Neurosci. 32, 12990–12998 (2012).

46. K. Umla-Runge, H. D. Zimmer, C. M. Krick, W. Reith, fMRI correlates of working memory: Specific posterior representation sites for motion and position information. Brain Res. 1382, 206–218 (2011).

47. T. B. Christophel, J. D. Haynes, Decoding complex flow-field patterns in visual working memory. Neuroimage 91, 43–51 (2014).

48. S. Liebe, G. M. Hoerzer, N. K. Logothetis, G. Rainer, Theta coupling between V4 and prefrontal cortex predicts visual short-term memory performance. Nat. Neurosci. 15, 456– 462 (2012).

49. S. J. D. Lawrence, et al., Laminar organization of working memory signals in human visual cortex. Curr. Biol. 28, 3435–3440 (2018).

50. T. Van Kerkoerle, M. W. Self, P. R. Roelfsema, Layer-specificity in the effects of attention and working memory on activity in primary visual cortex. Nat. Commun. 8, 13804 (2017).

51. Y. Merrikhi, et al., Spatial working memory alters the efficacy of input to visual cortex. Nat. Commun. 8, 15041 (2017).

52. R. L. Rademaker, C. Chunharas, J. T. Serences, Coexisting representations of sensory and mnemonic information in human visual cortex. Nat. Neurosci. 22, 1336–1344 (2019).

53. J. W. Bisley, D. Zaksas, J. A. Droll, T. Pasternak, Activity of neurons in cortical area MT during a memory for motion task. J. Neurophysiol. 91, 286–300 (2004).

54. M. Suzuki, J. Gottlieb, Distinct neural mechanisms of distractor suppression in the frontal and parietal lobe. Nat. Neurosci. 16, 98–104 (2013).

55. S. N. Jacob, A. Nieder, Complementary roles for primate frontal and parietal cortex in guarding working memory from distractor stimuli. Neuron 83, 226–237 (2014).

56. X. L. Qi, A. C. Elworthy, B. C. Lambert, C. Constantinidis, Representation of remembered stimuli and task information in the monkey dorsolateral prefrontal and posterior parietal cortex. J. Neurophysiol. 113, 44–57 (2015).

57. T. Haarmeier, T. Kammer, Effect of TMS on oculomotor behavior but not perceptual stability during smooth pursuit eye movements. Cereb. cortex 20, 2234–2243 (2010).

58. A. Zahorodnii, et al., Overcoming sensory-memory interference in working memory circuits. bioRxiv 2025.03.17.643652 (2025). 10.1101/2025.03.17.643652.

59. B. Lee, B. Pesaran, R. A. Andersen, Area MSTd neurons encode visual stimuli in eye coordinates during fixation and pursuit. J. Neurophysiol. 105, 60–68 (2010).

60. P. W. Dicke, S. Chakraborty, P. Thier, Neuronal correlates of perceptual stability during eye movements. Eur. J. Neurosci. 27, 991–1002 (2008).

61. W. O. Guldin, O. J. Grüsser, Is there a vestibular cortex? Trends Neurosci. 21, 254–259 (1998).

62. M. U. Trenner, et al., Human cortical areas involved in sustaining perceptual stability during smooth pursuit eye movements. Hum. Brain Mapp. 29, 300–311 (2008).

63. T. Haarmeier, P. Thier, M. Repnow, D. Petersen, False perception of motion in a patient who cannot compensate for eye movements. Nature 389, 849–852 (1997).

64. A. Lindner, T. Haarmeier, M. Erb, W. Grodd, P. Thier, Cerebrocerebellar circuits for the perceptual cancellation of eye-movement-induced retinal image motion. J. Cogn. Neurosci. 18, 1899–1912 (2006).

65. J. D. Murray, J. Jaramillo, X. J. Wang, Working memory and decision-making in a frontoparietal circuit model. J. Neurosci. 37, 12167–12186 (2017).

66. S. H. Lee, D. J. Kravitz, C. I. Baker, Goal-dependent dissociation of visual and prefrontal cortices during working memory. Nat. Neurosci. 16, 997–999 (2013).

67. T. B. Christophel, R. M. Cichy, M. N. Hebart, J. D. Haynes, Parietal and early visual cortices encode working memory content across mental transformations. Neuroimage 106, 198–206 (2015).

68. G. Pezzulo, T. Parr, K. Friston, Active inference as a theory of sentient behavior. Biol. Psychol. 186, 108741 (2024).

69. K. Matsuda, T. Nagami, Y. Sugase, A. Takemura, K. Kawano, A widely applicable real-time mono/binocular eye tracking system using a high frame-rate digital camera. Lect. Notes Comput. Sci. (including Subser. Lect. Notes Artif. Intell. Lect. Notes Bioinformatics) 10271, 593–608 (2017).

70. D. H. Brainard, The Psychophysics toolbox. Spat. Vis. 10, 433–436 (1997).

71. D. G. Pelli, The VideoToolbox software for visual psychophysics: Transforming numbers into movies. Spat. Vis. 10, 437–442 (1997).

72. M. Kleiner, et al., What’s new in psychtoolbox-3. Perception 36, 1–16 (2007).

73. T. Miyamoto, K. Numasawa, R. Hirano, Y. Yoshimura, S. Ono, Reduced latency in manual interception with anticipatory smooth eye movements. iScience 28, 111849 (2025).

74. T. Miyamoto, K. Numasawa, S. Ono, Changes in visual speed perception induced by anticipatory smooth eye movements. J. Neurophysiol. 127, 1198–1207 (2022).

75. E. Combrisson, K. Jerbi, Exceeding chance level by chance: The caveat of theoretical chance levels in brain signal classification and statistical assessment of decoding accuracy. J. Neurosci. Methods 250, 126–136 (2015).

