## Supplemental information for "Retinal image motion distracts visual motion memory even when generated by eye movement"

**Author Contributions:** Conceptualization, T.M., and K.N.; Methodology, T.M.; Software, T.M.; Formal Analysis, T.M.; Investigation, T.M., and K.N.; Writing – Original Draft, T.M.; Writing – Review & Editing, T.M. and K.N.; Funding Acquisition, T.M.

**Competing Interest Statement:** The authors declare no competing interests.

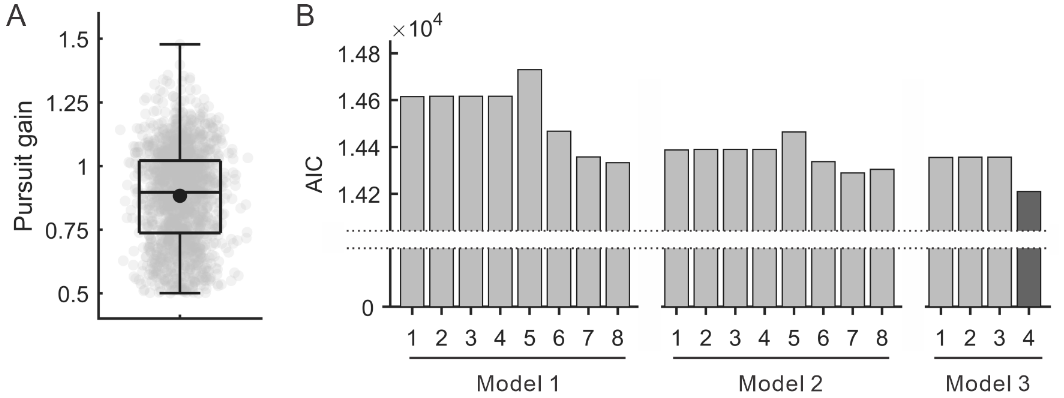


Model 1: response ~ 1 + condition * $\Delta$dist-speed$_{conditioned}$ * $\Delta$test-speed

1-1: [Linear predictor] + (1 | observer)
1-2: [Linear predictor] + (1 + condition | observer)
1-3: [Linear predictor] + (1 + $\Delta$dist-speed$_{conditioned}$ | observer)
1-4: [Linear predictor] + (1 + $\Delta$test-speed | observer)
1-5: [Linear predictor] + (1 + condition * $\Delta$dist-speed$_{conditioned}$ | observer)
1-6: [Linear predictor] + (1 + condition * $\Delta$test-speed | observer)
1-7: [Linear predictor] + (1 + $\Delta$dist-speed$_{conditioned}$ * $\Delta$test-speed | observer)
1-8: [Linear predictor] + (1 + condition * $\Delta$dist-speed$_{conditioned}$ * $\Delta$test-speed | observer)

Model 2: response ~ 1 + condition * $\Delta$dist-speed$_{actual}$ * $\Delta$test-speed

2-1: [Linear predictor] + (1 | observer)
2-2: [Linear predictor] + (1 + condition | observer)
2-3: [Linear predictor] + (1 + $\Delta$dist-speed$_{actual}$ | observer)
2-4: [Linear predictor] + (1 + $\Delta$test-speed | observer)
2-5: [Linear predictor] + (1 + condition * $\Delta$dist-speed$_{actual}$ | observer)
2-6: [Linear predictor] + (1 + condition * $\Delta$test-speed | observer)
2-7: [Linear predictor] + (1 + $\Delta$dist-speed$_{actual}$ * $\Delta$test-speed | observer)
2-8: [Linear predictor] + (1 + condition * $\Delta$dist-speed$_{actual}$ * $\Delta$test-speed | observer)

**Model 3: response ~ 1 + $\Delta$dist-speed$_{actual}$ * $\Delta$test-speed**

3-1: [Linear predictor] + (1 | observer)
3-2: [Linear predictor] + (1 + $\Delta$dist-speed$_{actual}$ | observer)
3-3: [Linear predictor] + (1 + $\Delta$test-speed | observer)
**3-4: [Linear predictor] + (1 + $\Delta$dist-speed$_{actual}$ * $\Delta$test-speed | observer)**

**Figure SI 1.** Supplementary data of Experiment 1

A. Distribution of pursuit gain for all 3,353 trials by 8 observers. Each dot corresponds to each trial, the box plot shows the interquartile range. The mean value was 0.88 (a black dot).

B. Variations in 20 GLMM sets. The selected model is shown as dark gray (Model 3-4).

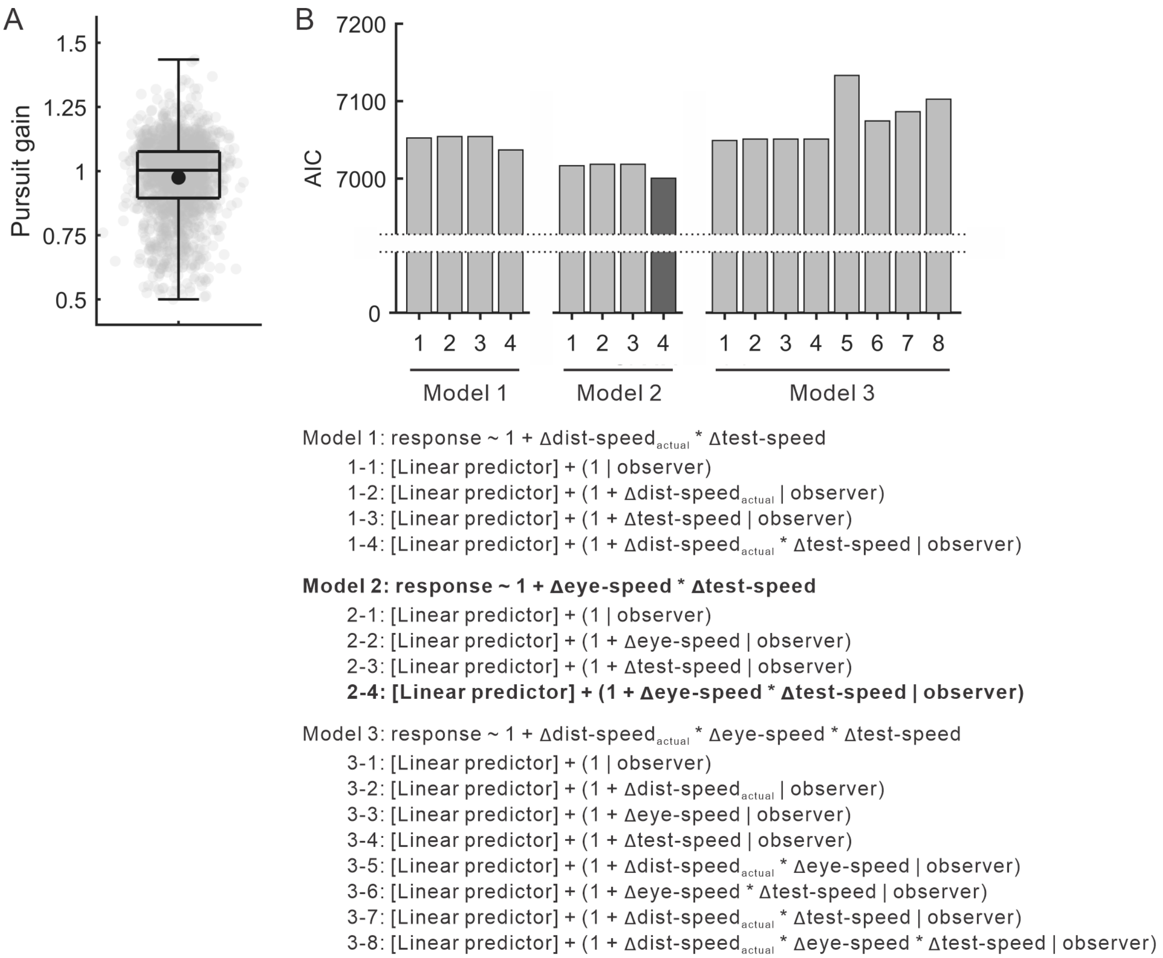


**Figure SI 2.** Supplementary data of Experiment 2

A. Distribution of pursuit gain for all 1,706 trials by 8 observers. Each dot corresponds to each trial, the box plot shows the interquartile range. The mean value was 0.97 (a black dot).

B. Variations in 16 GLMM sets. The selected model is shown as dark gray (Model 2-4).
